# Are some commercial diets inadequate in essential nutrients?

**DOI:** 10.1101/2021.03.31.437852

**Authors:** Alan M. Preston, Cindy A. Rodriguez, Marianna M. Preston

**Affiliations:** Department of Biochemistry, University of Puerto Rico, Medical Sciences Campus, San Juan, Puerto Rico, United States of America; NutriEtiquetas, Franklin, Massachusetts, United States of America

## Abstract

**Background:** Commercial manufacturers have formulated diets to promote not only weight reduction but also to reduce risks of chronic diseases. The objective of this study is to determine if these formulations satisfy requirements for essential nutrients.

**Methods:** We have selected two established commercial diets, one low fat, high carbohydrate (diet 1) and the other, high fat, low carbohydrate (diet 2) and determined “representative meals” through use of recipes suggested in the manufacturer’s manuals. Nutrition Data System for Research (NDSR) software has been used to perform the most extensive nutrient analysis to date of these diets. Tables report macronutrients (energy), vitamins, minerals, essential amino acids, essential lipids and nutrient-related components for a total of 62entries.

**Results:** Diet 1 satisfied requirements for 50 of these (81%) with only vitamin B12, vitamin D, and essential fatty acids not reaching recommended levels, while fiber and glycemic load exceeded suggested values. Diet 2 satisfied requirements for 46 of the components (71%) but had excess percentage of fat, especially saturated fat, sodium and cholesterol as well as decreased percentage of carbohydrate resulting in suboptimal intake of B-complex vitamins (B1, niacin and total folate) as well as fiber.

**Conclusions:** Neither diet satisfied adequacy for all reported nutrients. However, based on nutrient content alone diet 1, if supplemented or modified, could be sustained over the long term whereas diet 2 should not be encouraged for long term adaptation

## Introduction

In today’s obesogenic environment, more Americans are using some form of weight-reduction diet than were they ten years ago [1]. The good news is that weight reduction diets do “work” at least in the short term. A recent publication reported that all of 14 commercial diets ended up with weight loss at 6 months accompanied by favorable health results [2]. This pattern of short term weigh loss has been documented innumerable times by a multitude of investigators and summarized in several reviews [3-6], Nutrient composition of these diets varied widely among percent and types of carbohydrates, proteins, fats as well as micronutrients. The objective of our study is to determine if the manufacturers of commercial diets have formulated recipes which provided recommended dietary levels of essential nutrients. To accomplish this, we have chosen two popular commercial diets and obtained representative examples of each with use of suggested meal plans and determined nutrient adequacy with the use of software programs.

## Materials and methods

### Software

The dietary analysis is extensive using the full power of the Nutrition Data System for Research (NDSR) software of which the 2019 version contains 174 nutrients, nutrient ratios and other food components [7]. Nutrients having dietary reference index’s values (DRI’s) [8] or recommended dietary allowances (RDA’s) [9] will be reported. Other nutrients which can be biologically active but have no established recommendations such as phytochemicals found in plants in small amounts (polyphenolic flavonoids, carotenoids, etc. [10] and sugar alcohols, a class of polyols (sorbitol, mannitol, xylitol, etc.) which are present in varying levels in many fruits and vegetables [11] will not be reported. Likewise, non-essential amino acids and other non-essential nutrients found among the 174 entities in the NDSR will not be evaluated. Finally, some nutrient-related components (caffeine, glycemic load) will be reported making a total of 62 entries.

### Menus

We are using two established commercial diets, one is low fat, high carbohydrate, plant protein (Diet 1) [12]. This type of nutrient formulation is the basis for the Ornish, Macrobiotic and TLC diets, among others [13] although some incorporate animal protein. The other is high fat, low carbohydrate, mainly animal protein (Diet 2) [14]. This type of nutrient formulation is the basis for the Atkins, Paleo and Keto diets, among others. [13]. “Representative meals” were chosen through use of recipes suggested in the manufacturer’s manuals [12, 14]. We have selected, using a random numbers table, five of the 21 suggested daily menus from our designated commercial diet manuals which contain detailed content (ingredients and portion size) for breakfasts, lunches, dinners, and snacks. For clarification purposes, it should be noted that the diet manuals are structured differently. For diet 1, twenty one meal plans were listed in order (1 to 21). Therefore, the meal chosen by random numbers corresponds to the number in the list as shown in column 1, Table 1-top. For diet 2, twenty-one meal plans were listed but the order in which they were eaten was specified. (week and day). Therefore the meal chosen by random numbers corresponded to the specified week and day as shown in column 1, Table 1-bottom. Thumb-nail sketches of both diets are presented in table 1 to portray typical menus. Results of the five meals are averaged (representative meal) and reported in tables 2-6.

**Table 1.**
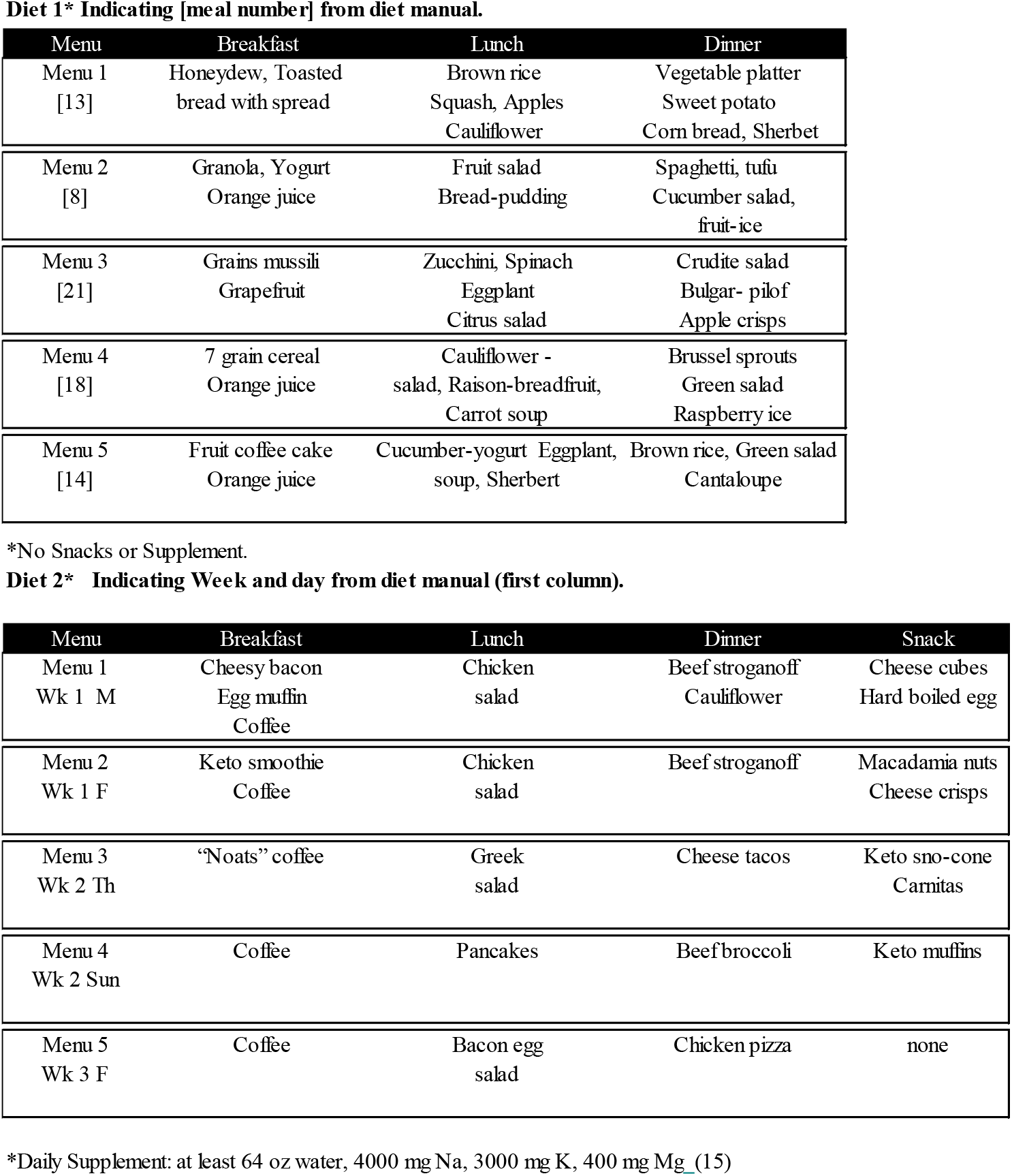
Menus for Diets

**Table 2.** Macronutrients

| Diet 1 |  |  |  |  |  |  | Diet 2 |  |  |  |  |  |  |
| --- | --- | --- | --- | --- | --- | --- | --- | --- | --- | --- | --- | --- | --- |
| Content (units) | Meal 1 | Meal 2 | Meal 3 | Meal 4 | Meal 5 | Mean ± SD | DRI | Meal 1 | Meal 2 | Meal 3 | Meal 4 | Meal 5 | Mean ± SD |
| Energy KCAL | 2,212.41 | 1,858.16 | 2,147.73 | 1,794.38 | 1,325.81 | 1867.7 ± 315.07 | 2000 | 1,736.49 | 2,288.84 | 1,404.57 | 900.65 | 845.31 | 1435.17 ± 539.28 |
| Total fat gm | 42.88 | 64.20 | 38.46 | 55.32 | 35.54 | 47.28 ± 10.82 | ≤70 | 132.74 | 161.69 | 115.87 | 74.78 | 70.15 | 111.05 ± 34.78 |
| Fat calories % | 17.44 | 31.10 | 16.12 | 27.74 | 24.13 | 23.31 ± 5.78 | 25-30 | 68.80 | 63.58 | 74.24 | 74.73 | 74.69 | 71.21 ± 4.42 |
| Total carbs gm | 406.46 | 300.67 | 396.20 | 295.50 | 238.46 | 327.46 ± 64.23 | 260 | 26.76 | 66.40 | 34.17 | 25.60 | 15.10 | 33.61 ± 17.49 |
| Carb calories % | 73.49 | 64.72 | 73.79 | 65.87 | 71.94 | 69.96 ± 3.88 | 45-65 | 6.16 | 11.60 | 9.73 | 11.37 | 7.15 | 9.2 ± 2.2 |
| Total prot gm | 87.12 | 45.33 | 71.92 | 59.64 | 37.18 | 60.24 ± 17.96 | 56 | 111.04 | 153.97 | 68.08 | 38.19 | 44.00 | 83.06 ± 43.76 |
| Prot calories % | 15.75 | 9.76 | 13.39 | 13.30 | 11.22 | 12.68 ± 2.05 | 10-35 | 25.58 | 26.91 | 19.39 | 16.96 | 20.82 | 21.93 ± 3.75 |
| Animal prot gm | 7.91 | 3.93 | 11.12 | 5.31 | 5.12 | 6.68 ± 2.57 | * | 104.06 | 135.03 | 46.40 | 29.82 | 35.41 | 70.14 ± 41.85 |
| Animal prot % | 9.08 | 8.68 | 15.46 | 8.91 | 13.76 | 11.18 ± 2.86 | * | 93.72 | 87.70 | 68.15 | 78.07 | 80.47 | 81.62 ± 8.7 |
| Vegetable prot gm | 79.21 | 41.40 | 60.80 | 54.33 | 32.07 | 53.56 ± 16.25 | * | 6.98 | 18.94 | 21.69 | 8.38 | 8.59 | 12.91 ± 6.13 |
| Vegetable prot % | 90.92 | 91.32 | 84.54 | 91.09 | 86.24 | 88.82 ± 2.86 | * | 6.28 | 12.30 | 31.85 | 21.93 | 19.53 | 18.38 ± 8.7 |
| Alcohol gm | 1.39 | 0.05 | 0.77 | 0.38 | 0.61 | 0.64 ± 0.45 | ‡ | - | - | 0.00 | 0.09 | - | 0.02 ± 0.04 |
| Alcohol cal% | 0.00 | 0.00 | 0.00 | 0.00 | 0.00 | 0 ± 0 | ‡ | 0.00 | 0.00 | 0.00 | 0.01 | 0.00 | 0 ± 0 |
| Total SFA gm | 5.94 | 5.58 | 3.79 | 4.89 | 3.44 | 4.73 ± 0.98 | ≤15 | 64.76 | 86.66 | 46.14 | 31.66 | 26.96 | 51.23 ± 22.07 |
| SFA cal % | 2.42 | 2.70 | 1.59 | 2.45 | 2.33 | 2.3 ± 0.38 | ≤7 | 33.57 | 34.07 | 29.56 | 31.63 | 28.70 | 31.51 ± 2.12 |
| Total MUFA gm | 8.63 | 9.72 | 6.16 | 8.20 | 6.03 | 7.75 ± 1.44 | ≤44 | 41.86 | 45.81 | 36.86 | 26.98 | 25.09 | 35.32 ± 8.12 |
| MUFA cal % | 3.51 | 4.71 | 2.58 | 4.11 | 4.09 | 3.8 ± 0.72 | ≤20 | 21.70 | 18.01 | 23.62 | 26.96 | 26.71 | 23.4 ± 3.33 |
| Total PUFA gm | 22.89 | 44.35 | 24.47 | 35.94 | 22.74 | 30.08 ± 8.66 | ≤22 | 13.49 | 14.62 | 23.40 | 9.59 | 12.67 | 14.75 ± 4.64 |
| PUFA cal % | 9.31 | 21.48 | 10.25 | 18.03 | 15.44 | 14.9 ± 4.61 | ≤10 | 6.99 | 5.75 | 14.99 | 9.58 | 13.48 | 10.16 ± 3.58 |
| Total Trans FA gm | 0.06 | 0.23 | 0.12 | 0.18 | 0.11 | 0.14 ± 0.06 | ≤2 | 3.24 | 3.08 | 2.11 | 1.37 | 0.93 | 2.14 ± 0.91 |
| Trans FA cal % | 0.03 | 0.11 | 0.05 | 0.09 | 0.07 | 0.07 ± 0.03 | ≤1 | 1.68 | 1.21 | 1.35 | 1.36 | 0.99 | 1.32 ± 0.23 |
| Total sugar gm | 42.19 | 23.45 | 31.05 | 26.61 | 17.46 | 28.15 ± 8.3 | 40 | 0.82 | 2.24 | 1.85 | 2.62 | 1.50 | 1.8 ± 0.62 |
| Added sugar gm | 22.96 | 5.86 | 4.16 | 14.32 | 6.02 | 10.66 ± 7.09 | 38 | 0.43 | 0.35 | 0.56 | 2.02 | 0.11 | 0.69 ± 0.68 |
| Total fiber gm | 63.83 | 45.69 | 48.05 | 59.22 | 42.48 | 51.85 ± 8.22 | 25-35 | 9.89 | 23.03 | 16.05 | 6.81 | 6.71 | 12.5 ± 6.27 |
| Water gm | 3,085 | 1,754 | 2,128 | 2,197 | 1,748 | 2182.6 ± 487.95 | 1811 P | 3,014 | 3,275 | 3,365 | 2,851 | 1,639 | 2828.75 ± 622.28 |

\* No DRI
‡ No recommendation dietary guideline for American (9) suggest moderate intake 2 drinks per day for Men / 1 drink per day for Women. (Drink = 12 oz beer or 5 oz wine)
P 8 Glasses x8 ounces = 64 oz of water (9)

### Statistics

A random numbers table (non-repeat) was used to select meals. Nutrient results from the NDSR software were recorded as meal content of five selected recipes from the manufacturer’s manuals for diets 1 and 2. The average and standard deviations for nutrients were calculated and compared to recommended guidelines.

## Results

A word on the manner of data presentation: When possible, we use RDA’s which are the daily dietary intake levels of nutrients considered sufficient by the Food and Nutrition Board of the Institute of Medicine to meet the requirements of 97.5% of healthy individuals in each life-stage and sex groups (10). Because of limited space, the RDA values listed will be for adult males; females have slightly lower values. In a few instances, reference values will be expresses as adequate intake (AI), defined as recommended average daily nutrient intake (9). Importantly, there is no RDA for energy (caloric intake) which depends on a myriad of individual factors. Consequently, energy and macronutrient content will be expressed as DRI values which give a rough idea of how much energy a person should be eating each day, and how much fat, sugar, salt and so on being based on an average-sized adult doing an average amount of physical activity. DRI values for energy have been set at 2000 Kcal for men and 1800 Kcal for women [9].

It can be seen that the sum of percentages of fat, carbohydrate and fat calories slightly exceeds the total Kcal in line 1, table 2 for both diets. This is due to the fact that calories from foods in the NDSR are determined chemically where energy values vary [19] whereas our calculations use standard values set for carbohydrates, protein and for fat of 4,4 and 9 Kcal/gm respectively.

There was moderate agreement in consistency of nutrient composition for most meals for diet 1, with a maximum difference of 900 Kcal between highest and lowest caloric ingestion, however, diet 2 had less agreement with a maximum difference of 9000 Kcal/gm. This caloric difference resulted in meal to meal variations of all other nutrients in the tables.

Of the 62 nutrients and nutrient- like components reported, fifty one (81%) achieved or fell within reference ranges for diet 1 and forty six (71%) for diet 2. Components outside reference ranges, both below and above include Diet 1: Table 2 (gm carbohydrate, %carbohydrate, fiber-all high), Table 3 (vitamin D, vitamin B12 –both low), Table 4 (sodium-low), Table 5 (essential fatty acids all low), Table 6 (cholesterol-low, glycemic load-high). For diet 2: Table 2 (gm fat and % fat, especially saturated fat-all too high, gm carbohydrate and % carbohydrate, fiber-all too low), Table 3 (Vitamin D, Vitamin B1, niacin, total folate-all too low, vitamin E-slightly low), Table 4 (sodium-too high), Table 5 (EPA, DHA-low), Table 6 (cholesterol-high).

**Table 3.**
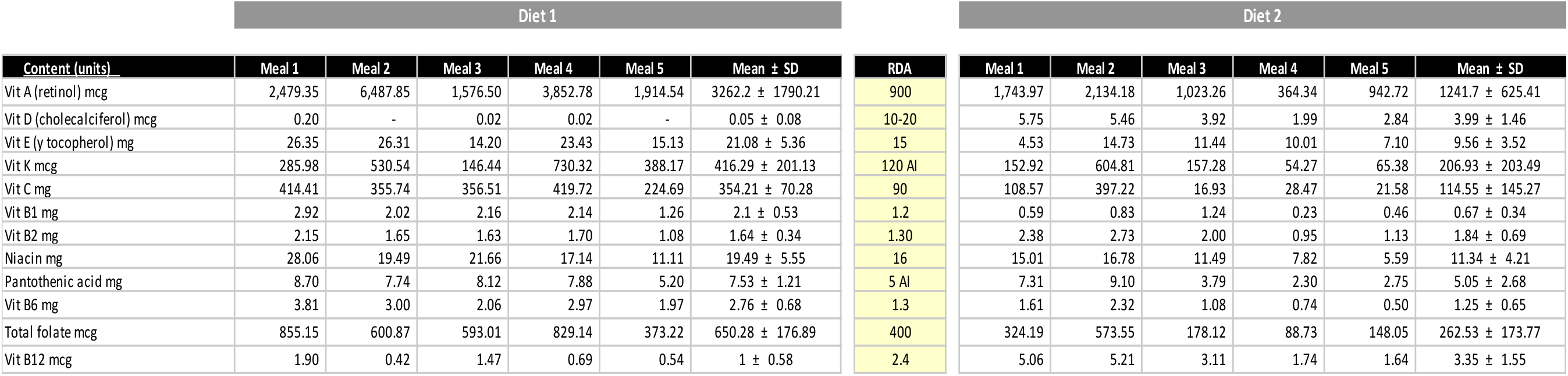
Vitamins

**Table 4.** Minerals

| Diet 1 |  |  |  |  |  |  | Diet 2 |  |  |  |  |  |  |
| --- | --- | --- | --- | --- | --- | --- | --- | --- | --- | --- | --- | --- | --- |
| Content (units) | Meal 1 | Meal 2 | Meal 3 | Meal 4 | Meal 5 | Mean ± SD | RDA | Meal 1 | Meal 2 | Meal 3 | Meal 4 | Meal 5 | Mean ± SD |
| Calcium mg | 1,275.19 | 604.36 | 841.54 | 794.88 | 578.56 | 818.91 ± 250.23 | 1000 | 760.26 | 1,818.96 | 1,164.94 | 377.51 | 472.28 | 918.79 ± 526.95 |
| Phosphorous mg | 2,126.66 | 1,159.89 | 1,485.69 | 1,349.59 | 1,026.99 | 1429.76 ± 382.17 | 700 | 1,134.73 | 1,764.05 | 1,504.68 | 530.56 | 763.95 | 1139.6 ± 455.04 |
| Magnesium mg | 796.39 | 491.30 | 497.90 | 506.37 | 423.58 | 543.11 ± 130.01 | 310 | 354.07 | 506.09 | 637.05 | 377.41 | 411.24 | 457.17 ± 103.78 |
| Iron mg | 23.19 | 13.56 | 13.73 | 14.02 | 11.52 | 15.21 ± 4.09 | 8 | 6.77 | 10.67 | 11.24 | 4.49 | 3.93 | 7.42 ± 3.04 |
| Zinc mg | 17.96 | 8.32 | 10.00 | 9.25 | 8.13 | 10.73 ± 3.68 | 11 | 11.37 | 14.70 | 12.31 | 6.26 | 4.90 | 9.91 ± 3.72 |
| Copper mg | 3.51 | 1.86 | 2.21 | 2.20 | 1.57 | 2.27 ± 0.66 | 0.9 | 0.67 | 1.25 | 1.54 | 0.71 | 0.87 | 1.01 ± 0.33 |
| Selenium mcg | 138.28 | 79.85 | 118.31 | 88.61 | 41.06 | 93.22 ± 33.41 | 55 | 132.67 | 133.96 | 78.57 | 44.07 | 596.68 | 197.19 ± 202.62 |
| Sodium mg | 440.53 | 384.99 | 399.06 | 442.86 | 211.24 | 375.74 ± 85.32 | 1500 | 4,914.97 | 7,342.25 | 4,041.47 | 5,132.50 | 5,034.59 | 5293.16 ± 1095.55 |
| Potassium mg | 7,461.92 | 5,065.32 | 4,433.99 | 5,005.98 | 4,085.53 | 5210.55 ± 1183.24 | 4700 | 4,117.21 | 5,593.91 | 4,083.22 | 3,274.73 | 3,184.26 | 4050.67 ± 864.85 |
| Manganese mg | 18.42 | 7.86 | 8.79 | 9.00 | 6.67 | 10.15 ± 4.22 | 1.8 | 0.79 | 1.93 | 3.98 | 1.32 | 0.91 | 1.79 ± 1.17 |

**Table 5.** Essential Amino Acids and Essential Fatty Acids

| Diet 1 |  |  |  |  |  |  | Diet 2 |  |  |  |  |  |  |
| --- | --- | --- | --- | --- | --- | --- | --- | --- | --- | --- | --- | --- | --- |
| Content (units) | Meal 1 | Meal 2 | Meal 3 | Meal 4 | Meal 5 | Mean ± SD | RDA | Meal 1 | Meal 2 | Meal 3 | Meal 4 | Meal 5 | Mean ± SD |
| Histidine mg | 2,010.0 | 1,034.0 | 1,583.0 | 1,364.0 | 812.0 | 1360.6 ± 419.2 | 14 | 2,607.0 | 3,757.0 | 2,099.0 | 1,108.0 | 1,282.0 | 2170.6 ± 962.4 |
| Isoleucine mg | 3,300.0 | 1,696.0 | 2,780.0 | 2,268.0 | 1,404.0 | 2289.6 ± 692.6 | 19 | 4,317.0 | 6,268.0 | 3,060.0 | 1,692.0 | 2,120.0 | 3491.4 ± 1655 |
| Leucine mg | 5,941.0 | 3,159.0 | 4,992.0 | 3,712.0 | 2,427.0 | 4046.2 ± 1265.3 | 42 | 7,999.0 | 11,748.0 | 5,299.0 | 2,989.0 | 3,606.0 | 6328.2 ± 3218.3 |
| Lysine mg | 4,039.0 | 2,243.0 | 2,901.0 | 2,916.0 | 1,914.0 | 2802.6 ± 728.6 | 38 | 7,330.0 | 10,950.0 | 4,267.0 | 2,694.0 | 3,121.0 | 5672.4 ± 3097 |
| Methionine mg | 1,296.0 | 758.0 | 1,206.0 | 833.0 | 585.0 | 935.6 ± 271.3 | 19 | 2,354.0 | 3,144.0 | 1,599.0 | 896.0 | 1,365.0 | 1871.6 ± 791.8 |
| Phenylalanine mg | 3,826.0 | 2,138.0 | 3,402.0 | 2,475.0 | 1,628.0 | 2693.8 ± 809.5 | 33.0 | 4,416.0 | 6,168.0 | 3,025.0 | 1,705.0 | 2,075.0 | 3477.8 ± 1639.1 |
| Tryptophane mg | 987.0 | 550.0 | 762.0 | 654.0 | 433.0 | 677.2 ± 189.5 | 20 | 1,052.0 | 1,764.0 | 987.0 | 326.0 | 548.0 | 935.4 ± 494.8 |
| Tyrosine mg | 2,605.0 | 1,286.0 | 2,051.0 | 1,585.0 | 1,116.0 | 1728.6 ± 541.1 | 5 | 3,622.0 | 5,068.0 | 2,535.0 | 1,241.0 | 1,715.0 | 2836.2 ± 1377.7 |
| Valine mg | 4,037.0 | 2,284.0 | 3,488.0 | 2,848.0 | 1,878.0 | 2907 ± 782.9 | 24 | 5,401.0 | 7,763.0 | 3,672.0 | 1,915.0 | 2,405.0 | 4231.2 ± 2138 |
| Alpha linolenic acid (ALA) mg | 1,261.0 | 1,101.0 | 561.0 | 728.0 | 640.0 | 858.2 ± 273.5 | 1600 | 1,522.0 | 1,909.0 | 4,809.0 | 271.0 | 277.0 | 1757.6 ± 1660.8 |
| Eicosapantanaeolic acid (EPA) mg | - | - | - | - | - | 0 ± 0 | † | 27.0 | 20.0 | 14.0 | 8.0 | 12.0 | 16.2 ± 6.6 |
| Docosahexaeonic acid (DHA) mg | - | - | - | - | - | 0 ± 0 | † | 60.0 | 37.0 | 2.0 | 17.0 | 44.0 | 32 ± 20.4 |

\*Arginine is conditional † EPA + DHA = 250-500 mg/day (17)

**Table 6.** Nutrient-Related Components

| Diet 1 |  |  |  |  |  |  | Diet 2 |  |  |  |  |  |  |
| --- | --- | --- | --- | --- | --- | --- | --- | --- | --- | --- | --- | --- | --- |
| Content (units) | Meal 1 | Meal 2 | Meal 3 | Meal 4 | Meal 5 | Mean ± SD | DRI | Meal 1 | Meal 2 | Meal 3 | Meal 4 | Meal 5 | Mean ± SD |
| Cholesterol mg | 2.74 | 1.37 | 3.88 | 1.84 | 1.78 | 2.32 ± 0.9 | ≤ 300 mg | 934.13 | 712.80 | 256.86 | 296.71 | 477.77 | 535.65 ± 256.26 |
| Caffeine mg | - | - | - | - | - | 0 ± 0 | * | 189.44 | 100.90 | 189.53 | 94.81 | 94.72 | 133.88 ± 45.46 |
| Glycemic load | 171.72 | 144.54 | 177.28 | 103.86 | 92.80 | 138.04 ± 34.44 | † | 7.16 | 17.80 | 7.13 | 6.00 | 2.61 | 8.14 ± 5.11 |

\* ≤ 400 mg (16) † Glycemic Load (or GL) combines both the quantity and quality of carbohydrates . Low GL is between 1 and 10; a moderate GL is 11 to 19; and a high GL is 20 or higher (18).

The following discussion section will include comments on the enumeration of these outliers.

## Discussion

Diet manufacturers often shuffle proportions and types of carbohydrates, fats and proteins to create eating plans concomitant with reducing risk of major degenerative diseases commonly found in the United States [13]. Diet 1 having low fat, high carbohydrate and plant-based protein, which the manufacturer refers to as “heart friendly”, incorporates nutrients associated with favorable cardiovascular function [20]. These include high fiber, no animal protein as well as low fat (especially saturated fat), low sodium and cholesterol and little added sugar. As a consequence, the aberrant values in the tables for fiber, sodium and cholesterol are a result of conscious action of the manufacturer’s design of the diet. One of the most consistent results in table 2 is percent carbohydrate (about 70%) which is probably the most important ingredient in the diet’s formulation and used as a set point. In doing this, other components may be left short of achieving reference values. The absence of animal protein would account for low vitamin B12 which is exclusively formed in animals and probably the low vitamin D result since dairy products, a principle source of this vitamin, are minimized. The low amount of fat could account for diminished levels of alpha linolenic acid, especially for EPA and DHA (which measured zero) as well as for vitamin D. Finally, there is the issue of high glycemic load. Whether this is a matter of concern depends on the form of carbohydrate present [18]. Referring to Table 2, added sugar which has unfavorable circumstances on blood sugar is twofold below levels associated with risk while fiber which mitigates the rise in blood sugar is twice the recommended level fhus concern of high glycemic load should be minimized.

Diet 2 which is high fat, low carbohydrate and moderate protein is claimed by the manufacturer to be “fat burning” since very low carbohydrate intake (20 -35 gm/day at the start of the diet) triggers mobilization of lipid stores stimulating formation of ketone bodies which can have beside weight loss, therapeutic benefits such as reducing risk of insulin resistance and type 2 diabetes [21].

It should be mentioned at this point that our paper solely evaluates nutrient adequacy of the two diets and makes no judgment of manufacturer’s health claims. Diet 2 has been formulated to promote ketogenic metabolism [14]. This is accomplished by high fat content and very low amount of carbohydrate. Consequently, the aberrations in the results section for total fat, percent fat, total saturated fat, percent saturated fat, total carbohydrates and percent carbohydrates are intentionally made by the manufacturer. End results of this formulation are high dietary cholesterol and diminished intake of B-complex vitamins (Vitamin B1, niacin, total folate) and fiber which are all associated with carbohydrate content. An additional effect of low amount of carbohydrate is loss of body water. To prevent dehydration and electrolyte imbalance, at least eight glasses of water, 8 oz each are recommended accompanied by at least 4000 mg sodium (well above the DRI), 3000 mg potassium and 400 mg magnesium [15].. Since the amount of fat and animal protein are in abundance, one would not expect reduced levels of fat soluble vitamins or essential fatty acids as reported in tables 3 and 5.. A possible explanation is that the 5 meals selected missed menus that included seafood products of which several were included in the recipe manual.

Strengths of this study include the manner of data entry and diet analysis. Many dietary studies examine the amount of nutrients consumed by individuals which is susceptible to recall errors. Here we have exact ingredients and portion sizes, copied directly from the recipe books. In addition, using the full capacity of the NDSR software, we are able to perform the most extensive analyses of commercial weight loss diets to date.

Potential weaknesses include estimated intake of minor nutrients and determination of a “representative meal.” The number of days to validate intake of nutrients has been established using food frequency questionnaires (ffq’s), the results of which vary widely. Macronutrients (found in table 2) can be validated within a week, while some micronutrients (tables 3 and 4) may take a month or more [22]. In regard to “reference meal”, determination was made using the average of 5 meals, selected at random from the 21 daily meals in the diet manuals. Even though recipes are formulated to produce a relatively consistent meal content, there is still a variation of 900 Kcal between highest and lowest meal energies for diet 1 and 9000 Kcal for diet 2. A “true meal” would require analysis of all 21 meals in the diet manual, however, we believe this result would not differ substantially from our estimate and conclusions remain valid. Although as mentioned previously in the results section, all reference values in the tables are based solely on those of adult men and would not necessarily apply for women or children.

Finally, returning to the question posed in the title of this manuscript: “Are some commercial diets inadequate in essential nutrients?” The answer (in this case) is “yes” For diet 1, the formulation of macronutrients resulted in suboptimal ingestion of animal protein causing a deficiency of vitamin B12 and vitamin D, and the low fat also restricted vitamin D intake as well as reducing essential fatty acid content. The high level of fiber could furthermore compromise absorption of minerals [23]. Overall, diet 1 or eating patterns of similar composition should be “safe” over a long term if accompanied with a vitamin/mineral/essential fatty acid supplement or if modified from only plant protein to one incorporating some meat and seafood. For diet 2 the formulation of macronutrients resulted in excess amount of fat and fat associated nutrients as well as an insufficiency of carbohydrate and carbohydrate associated nutrients. To comply with DRI/RDA recommendations the formulation of diet 2 would have to be modified, reducing fat and increasing carbohydrate. This alteration however would defeat the ketogenic metabolic scheme and its purpose. Overall, diet 2 or eating patterns of a similar composition would be unsafe over the long term.

## Conclusions

Although the two commercial weight reduction diets we have chosen differ greatly in composition and have been formulated to promote dissimilar modes of action to reducing risk for chronic diseases, they both satisfy recommendations for most nutrients, being 81% for diet 1 and 71% for diet 2. The manner in which they differ is that diet 1 is sustainable over time if supplemented or modified whereas diet 2 is not sustainable over time due to nutritional imbalances and should not be continued.

## Acknowledgments

The authors would like to thank Cristina Palacios, PhD - Florida International University, Miami, FL allowing use the NDSR program as part of her grant number 1R01HD098589-01. Permission to use her name was obtained.

## Author Contributions

Alan M Preston contributed with: Conceptualization, Methodology, Validation, Formal Analysis, Investigation; Resources, Data curation; and Writing – Original Draft Preparation.

Cindy A Rodriguez contributed with: Software, Validation and Formal Analysis.

Marianna M Preston contributed with: Software, Validation, Formal Analysis and Data Curation.

All authors critically reviewed and approved the final version of the manuscript submitted for publication.

